# Fast Variational Alignment of non-flat 1D Displacements for Applications in Neuroimaging

**DOI:** 10.1101/2020.06.27.151522

**Authors:** Philipp Flotho, David Thinnes, Bernd Kuhn, Christopher J. Roome, Jonas F. Vibell, Daniel J. Strauss

**Affiliations:** Systems Neuroscience and Neurotechnology Unit, Neurocenter, Faculty of Medicine, Saarland University, Homburg/Saar, Germany; Summer Program, Japan Society for the Promotion of Science (JSPS), Tokyo, Japan; Department of Psychology, University of Hawai’i at Mānoa; Optical Neuroimaging Unit, Okinawa Institute of Science and Technology Graduate University, Tancha, Onna-son, Kunigami, Okinawa, Japan

**Keywords:** 1D alignment, 1D displacement, Variational methods, Evoked potentials, Event related potentials, Line scan, Two-photon microscopy, Confocal microscopy

## Abstract

**Background:** In the context of signal analysis and pattern matching, alignment of 1D signals for the comparison of signal morphologies is an important problem. For image processing and computer vision, 2D *optical flow* (OF) methods find wide application for motion analysis and image registration and variational OF methods have been continuously improved over the past decades.

**New Method:** We propose a variational method for the alignment and displacement estimation of 1D signals. We pose the estimation of non-flat displacements as an optimization problem with a similarity and smoothness term similar to variational OF estimation. To this end, we can make use of efficient optimization strategies that allow real-time applications on consumer grade hardware.

**Results:** We apply our method to two applications from functional neuroimaging: The alignment of 2-photon imaging line scan recordings and the denoising of evoked and event-related potentials in single trial matrices. We can report state of the art results in terms of alignment quality and computing speeds.

**Existing Methods:** Existing methods for 1D alignment target mostly constant displacements, do not allow native subsample precision or precise control over regularization or are slower than the proposed method.

**Conclusions:** Our method is implemented as a MATLAB toolbox and is online available. It is suitable for 1D alignment problems, where high accuracy and high speed is needed and non-constant displacements occur.

## 1 Introduction

*Optical flow* (OF) estimation methods form an important family of image processing algorithms with applications in many areas where motion analysis, tracking or registration is needed. In the context of OF estimation of 2D natural images and image registration, *variational methods* form a powerful group of well-studied methods that perform with very high accuracy in the presence of small displacements [1]. Up until today, they find applications in state of the art OF algorithms, where they can be used for the refinement of small displacements [2] or for the generation of ground truth displacement fields [3], [4]. They are able to estimate non-parametric, dense and smooth displacement fields with subpixel accuracy [1].

The displacement field is calculated by minimizing an energy functional with a data term as the error measure and a regularization term that enforces smooth solutions. The data term and regularizer are modeled with respect to the data characteristics and the expected type of motion.

Most commonly, variational OF methods are solved within an Euler-Lagrange framework, where a linearized approximation of the problem is solved with a coarse-to-fine strategy. This yields a sequence of convex programming problems, that can be optimized efficiently with parallel implementations on graphics processors [5], [6], [7].

Imaging areas, that require very high frame rates or spectral rather than spatial resolutions are suitable for the application of line scan sensors. Line scans are sequences of 1D recordings typically of a scene under static conditions or of a scene with camera or object movement orthogonal to the spatial dimension. Among others, they find application in industrial quality inspection [8] and biomedical imaging [9]. There are many sources for movement artifacts in line scan recordings: In the context of moving objects, poor synchronization, or jitter in the object movement results in non-uniform sampling. Brosch et al. [8] have proposed a variational formulation to estimate the warping function in the context of movement jitter for light field line scan data to reconstruct a recording that is sampled at even intervals.

Movements parallel to the spatial dimension result in shifts of the signal, that can often be approximated by constant displacements. Depending on the camera and lens / illumination setup, movements towards the camera can have additional effects such as linear displacements due to zooming or changes in signal morphology.

Within the scope of single trial analysis of *evoked and event-related potentials* (EP and ERP) in *electroencephalography* (EEG) recordings, event locked responses are analyzed [10]. The average response may have a high *signal to noise ratio* (SNR) but also a lower resolution of the signal. From the average signal it is also not possible to analyze temporal variations across trials. Popular approaches to increase SNR in ERP recordings for single trial analysis are often motivated by image processing methods [11]. This is because the single trial matrix representation of ERP recordings resembles the 2D representation of line scan data: Each line is a measurement with high temporal resolution of a stimulus locked event.

To remove constant displacements in 1D data, methods that maximize the crosscorrelation with respect to a reference profile find wide application. Methods to compare two arbitrary 1D signals are often based on *dynamic time warping* (DTW) [12]. The idea of DTW is to repeat individual samples of both signals to minimize a distance metric. This way, distinct features of the two signals can be directly compared. Nielsen et al. [13] introduced a method where they extend DTW with piecewise cross-correlation. Their correlation optimized warping approach is applied for the alignment of chromatographic data [14], [15].

In their preliminary work, Deriso et al. [16] have presented a variational formulation of the DTW problem. In their method they optimize the complete non-parametric warping function. They solve the non-convex optimization problem with dynamic programming in a multigrid framework.

In this work, we investigate non-parametric displacement estimation for 1D data and describe a variational method for non-parametric, dense displacement estimation of 1D signals, which is motivated by variational OF estimation.

By exploiting the linear motion assumption within a coarse-to-fine strategy, our approach solves a sequence of convex problems. We demonstrate the performance of our algorithm on two problems from neuroimaging with alignment of optical imaging line scan recordings and the compensation of latency jitter in ERP recordings. Our implementation is fast enough to allow real time alignments on consumer grade hardware.

## 2 Materials and Methods

In this section we describe our method for the alignment and displacement analysis of 1D signals. To this end, we propose a variational model and pre-processing strategy and describe our implementation.

We work with 2D representations of multiple 1D signals which can be temporal (EP and ERP alignment) or spatial (line scan alignment). Let *m* be the length (spatial or temporal) of the 1D signal and *n* be the number of measurements. Then an individual measurement is given by 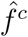: Ω_1_ → ℝ with *x* ∈ Ω_1_ = (0, *m*) ⊂ ℝ. The 2D matrix representation of multiple measurements is given by the function *f* ^*c*^(*x, y*): Ω_2_ × ℕ → ℝ, where (*x, y*) ∈ Ω_2_ = (0, *m*) × (0, *n*) ⊂ ℝ^2^ are the coordinates. For multichannel recordings *c* = 1, *…, k* ⊂ ℕ denotes the discrete number of the channel. The first dimension is either spatial or temporal and goes along the 1D signals and the second dimension is orthogonal to the signal and usually temporal and represents individual measurements. We depict the reference profile with *g*^*c*^(*x*): (0, *m*) → ℝ. The discrete version of *g* is denoted by 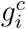 and of *f* by 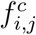 where the dimension along the first dimension is sampled uniformely and along the second can be arbitrarily sampled and explicitly defined by the input data. By means of simplicity, we depict partial derivatives with lower case letters *x* and *y* and use the lower case letters *i, j* exclusively for discretized access. Continuous access to the discrete data is realized via bicubic interpolation as recommended by [1].

### 2.1 Variational Model for 1D Alignment

We formulate the problem of aligning 1D signals as a variational problem, where we want to find displacements *u*(*x, y*): Ω_2_ → ℝ of 1D signals 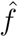 with respect to a reference *ĝ*. We assume gradient constancy between 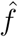 and *ĝ* as well as small gradients in *u*. We penalize both assumptions with the generalized Charbonnier penalty function Ψ_*a*_. Let *a* > 0, then the generalized Charbonnier penalty function [1] is given by

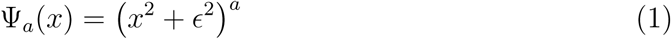

Following the authors in [1], we choose the penalization of the data and smoothness terms with *a*_*s*_, *a*_*d*_ ∈ [0.45, 1] ⊂ ℝ, where Ψ_0.5_ gives the Chabonnier penalty function and Ψ_1_ is quadratic penalization. Ψ_0.45_ is slightly non-convex but has been shown to perform well in the context of OF estimation [1]. The energy functional for 1D displacements of single channel signals is then given by

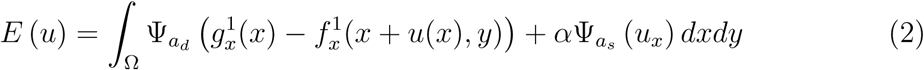

where the relation of the data and smoothness term is weighted by the smoothness weight parameter *α* ∈ ℝ_>0_. The displacement field *u* is then calculated as argmin *E* (*u*). To achieve that, we linearize the constancy assumption into *u*·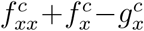 on a Gaussian pyramid and calculate the increments on each level.

The linearized assumption is a good approximation for small displacements and it has been shown, that the final solution of such a scheme approximates the unlinearized constancy assumption [6]. Gradient constancy with subquadratic penalization is invariant under global additive changes of the signal and robust with respect to outliers. We normalize the input data, which can additionally compensate for multiplicative changes. In the context of variational multimodal image registration, there exist intensity similarity measures [17], that allow affine (cross correlation), functional (correlation ratio), and arbitrary (mutual information) dependence of the two signals. Cross correlation is a commonly used metric for 1D signal analysis. Additionally to the gradient constancy assumption, we implement a cross correlation data term as presented in [17] with isotropic smoothness term, however the gradient descent scheme from [17] has much slower convergence than our gradient constancy model.

The variational formulation allows a straightforward extension to data with multiple channels. Given data *f* ^*k*^ with k channels, we can extend our data term with separate penalization on each channel as in

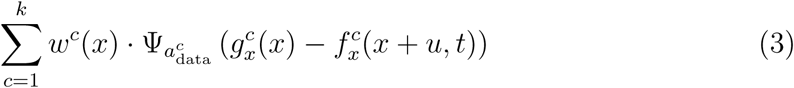

The weight term *w*(*x*): Ω → ℝ is multiplied pointwise to each channel and can be used to integrate knowledge on the information content / SNR per channel and to change the weight of each channel or datapoint.

To our knowledge, there exist no methods for the alignment of 1D signals that natively support the integration of multiple channels to calculate displacements.

#### Pre-Processing

In the following, we apply 1D Gaussian convolution with a fixed standard deviation 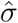 along the first dimension before calculating the displacements. This reduces high frequency noise and makes the signals differentiable. We normalize the input data to compensate for different scaling.

On top of that, due to potentially low SNR of the input data, we need an additional pre-processing strategy. We propose a novel, anisotropic kernel for the processing of 2D single trial matrices (see figure 1).

**Figure 1:**
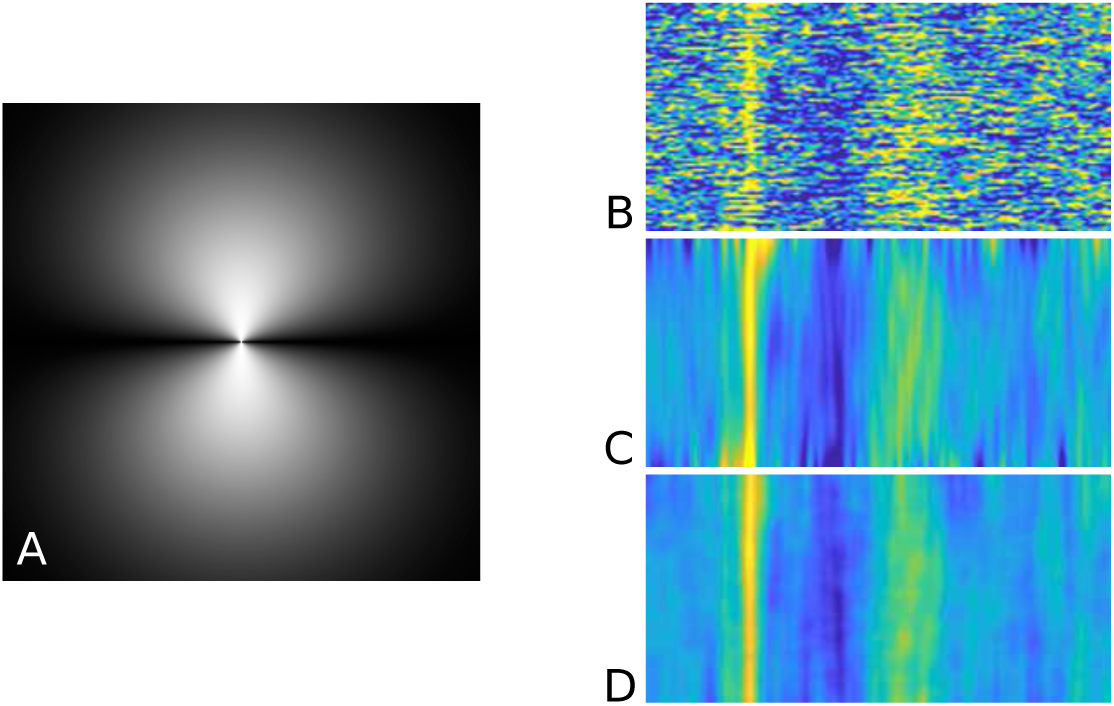
The anisotropic kernel (A) defined by equation 4 which we use for pre– smoothing: A normalized Gaussian kernel *K*_*σ*_ with zero mean and standard deviation *σ* that is weighted by the spatial angle. Applied on an ERP map (B), we see that traces across trials are accentuated (D) when comparing with Gaussian convolution (C). For (C) and (D) we used a standard deviation of *σ* = 10.

We choose a filter that smooths orthogonally to the discrete time sequences and line scans, while allowing displacements of the individual measurements. Our kernel is a 2D Gaussian function *G*_*σ*_ with zero mean and standard deviation *σ* = (*σ*_1_, *σ*_2_)^⊤^

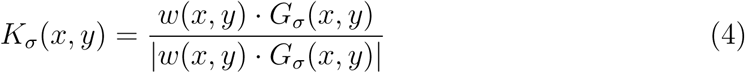

that is weighted by the angular distance to the 1D signals

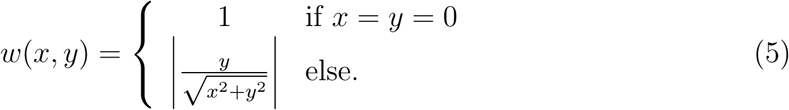

For the rest of the paper we use equal standard deviation *σ*_1_ = *σ*_2_.

In the context of real time alignment of data with low SNR, at each point in time we have only access to the past measurements. In this case, we can use half the kernel and weight *G*_*σ*_ with

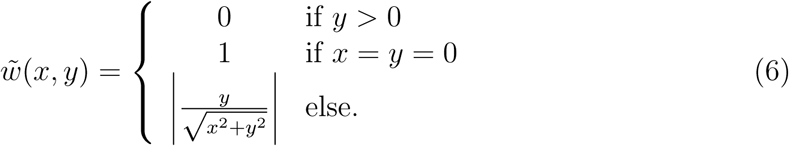

### 2.2 Implementation

We discretize the Euler-Lagrange equations of the linearized functional following Brox et al. [6] and solve it with an iterative solver with lagged diffusivity on a Gaussian scale space. It has been empirically shown, that median filtering of 2D flow increments improves the estimation accuracy [1]. Accordingly, at each pyramid level, we apply a 1D median filter to the estimated displacement increments. For the results presented in this paper, we choose a filtersize of 5. We use cubic interpolation for the compensation of displacement increments as well as for the final alignment. For multimodal data terms, we follow Hermosillo et al. [17] and implement a gradient descent algorithm on a Gaussian scale space with cross-correlation in the data term.

Our model requires a common reference and computes displacements and alignment for all measurements with respect to this reference. Analysis of the displacements of subsequent measurements can be realized from the displacement field without explicitly comparing two signals. However, for certain data, registration to a fixed reference can result in large displacements. Large displacements can violate the linearized constancy assumption on the highest level of the pyramid and hinder the estimation of the joint pdf for the multimodal data terms.

For this reason, we assume that we can find a row order with gradually increasing displacement magnitudes, meaning for all *i* ∈ 1, *…, m* we want to find the bijective function *π*(*x*): ℕ → ℕ such that the displacement magnitudes are gradually increasing |*u*_*i,π*(1)_| ≤ |*u*_*i,π*(*j*)_| ≤ |*u*_*i,π*(*n*)_| with respect to the reference profile. This mapping *π* integrates experiment specific a-priori knowledge into the alignment calculation or can be estimated heuristically. For example, we expect line scan data with high temporal resolution to show small inter line movements making the identity a reasonable choice for *π*.

In terms of implementation, for all *i* ∈ 1, *…, m* we initialize *u*_*i,π*(1)_ with a constant displacement field and then calculate the displacements row by row, where we initialize the highest pyramid level of row *π*_*i,j*_ with the previous result *u*_*i,π*(*j*−1)_. This strategy results in the smallest possible displacement increments, as long as we can provide some good *π*.

The parameters of our method are on the one hand the solver specific parameters such as warping depth, number of iterations, update frequency of the non-linear term and the warping stepsize *η* which can be increased for better accuracy. On the other hand, we have model specific parameters which modify the objective function: The smoothness parameter *α* defines the weight of the smoothness term with respect to the data term. The penalizer exponents of the generalized Charbonnier function 0.45 ≤ *a*_*d*_, *a*_*s*_ ≤ 1 control how deviations from the model assumptions are penalized: Large values of *a*_*s*_ enforce smooth displacements and low values allow discontinuities. Smaller values of *a*_*d*_ make the data term more robust with respect to outliers and noise.

We have implemened our method in MATLAB with the MATLAB Image Processing Toolbox and C++. The warping loop is written in MATLAB and on grounds of computing speed, the iterative solver is written in C++ and compiled into a `mex` function.

## 3 Results

In this section, we report the performance of our alignment approach on EEG single trial matrices and functional 2-photon imaging line scan recordings. We use real physiological as well as synthetic data. For the sake of visualization, we report the results for line scans (section 3.2) always transposed, meaning the displacements have been estimated along the y-axis. For quantitative analysis, we compare variational 1D alignment with the removal of constant shifts by maximizing cross correlation (no subsample refinement) and correlation optimized warping as implemented in [15].

### 3.1 Synthetic Experiments

We use two different synthetic datasets: The first one simulates a line scan recording with a non-constant motion artifact, that is difficult to remove. We use it to demonstrate the compensation by our method of motion artifacts that cannot be parametrized by constant shifts. To investigate overfitting and clustering of signal features as well as to benchmark computing speed, we use random 2D matrices.

#### Line Scan with Diverging Structure

For our first synthetic data experiment, we simulate a line scan recording that has been contaminated by non-constant motion noise. The data consists of 800 line measurements with 150 samples each. The average of the ground truth displacement field *û* is zero, which means the motion cannot be parametrized by constant shifts. The signal features converge and diverge in the center as seen by changes in *û*_*x*_ (see figure 2).

**Figure 2:**
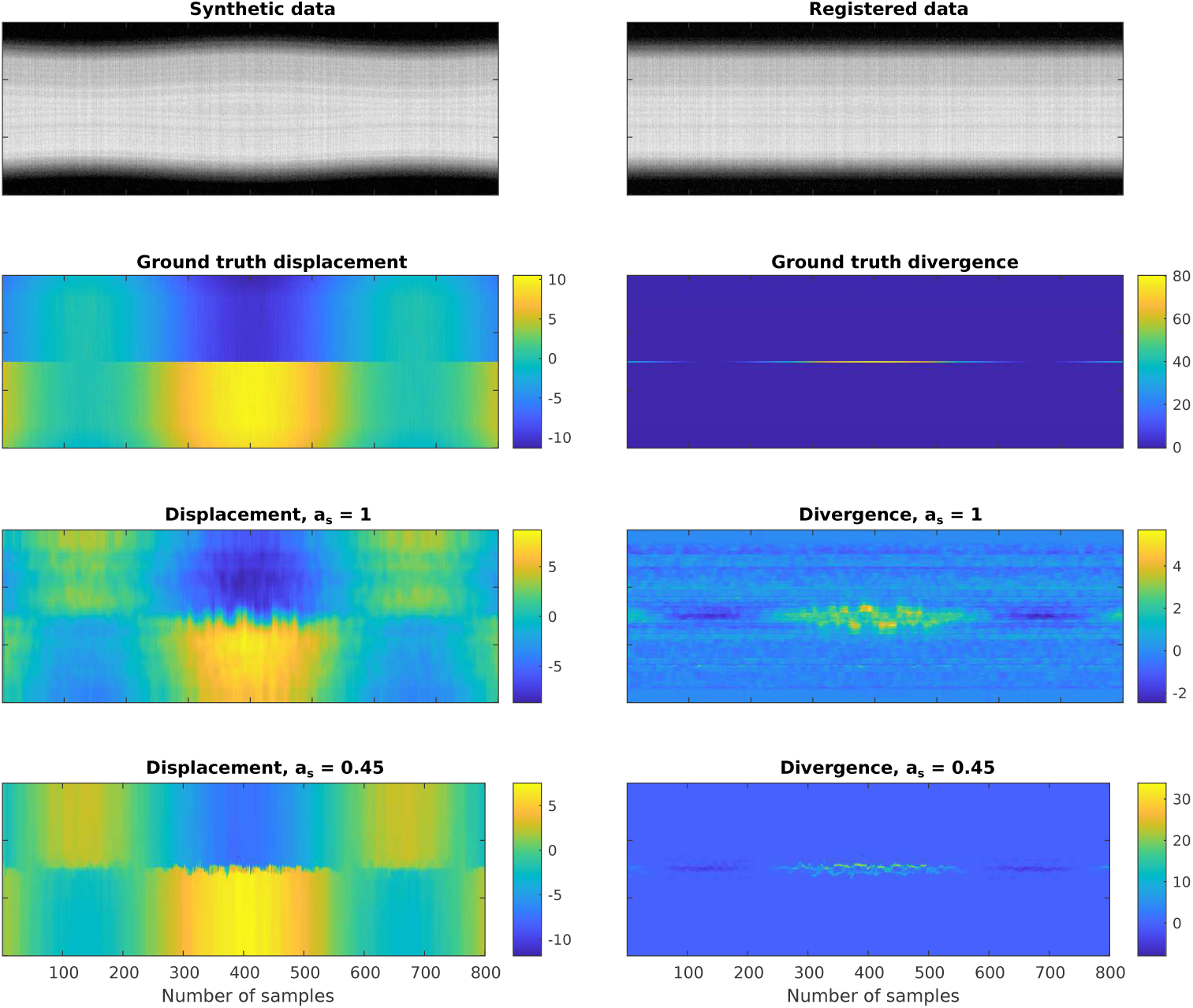
Synthetic line scan recording with converging and diverging features (top left) which can be almost completely compensated by our method (top right). The bottom left plots depict the ground truth displacement map and the estimated displacement map with *a*_*s*_ = 1 and *a*_*s*_ = 0.45. On the right is the first derivative in along the spatial dimension of the displacement map. The ground truth displacement map contains discontinuities which is why *a*_*s*_ = 0.45 yields a better estimation than *a*_*s*_ = 1.

For the evaluation, we use different values of *a*_*s*_ and choose parameters with less iterations for a fast estimation (see table 1). We choose the average profile as reference, which in the case of the variational method is refined one time. As expected, the dataset cannot be denoised with constant displacements (see figure 3 and table 1). Constant alignment with maximization of the cross correlation does not improve the signal. Quantitatively, correlation optimized warping performs even worse than constant alignment, which might be due to the high amount of noise in our data set. Because of the discontinuities in the displacement fields, a low choice of *a*_*s*_ performs better than quadratic penalization of the smoothness.

**Table 1:** Performance of our method in comparison with cross correlation based constant alignment (no subsample refinement) and correlation optimized warping. Smaller values for the average *standard deviation* (STD) and larger values for the average *peak signal to noise ratio* (PSNR) are better. *Variational fast* uses less iterations, smaller *η* and no refinement of the reference.

| Method | STD | PSNR | Time (ms) |
| --- | --- | --- | --- |
| Correlation optimized warping | 0.048 | 26.09 | 58.5 |
| Raw signal | 0.046 | 26.51 | — |
| Constant displacement cross correlation | 0.046 | 26.52 | 0.1 |
| Variational fast $a_s = 0.45$ | 0.034 | 29.25 | 1.2 |
| Variational $a_s = 1$ | 0.033 | 29.43 | 7.8 |
| Variational $a_s = 0.45$ | 0.033 | 29.55 | 9.3 |

**Figure 3:**
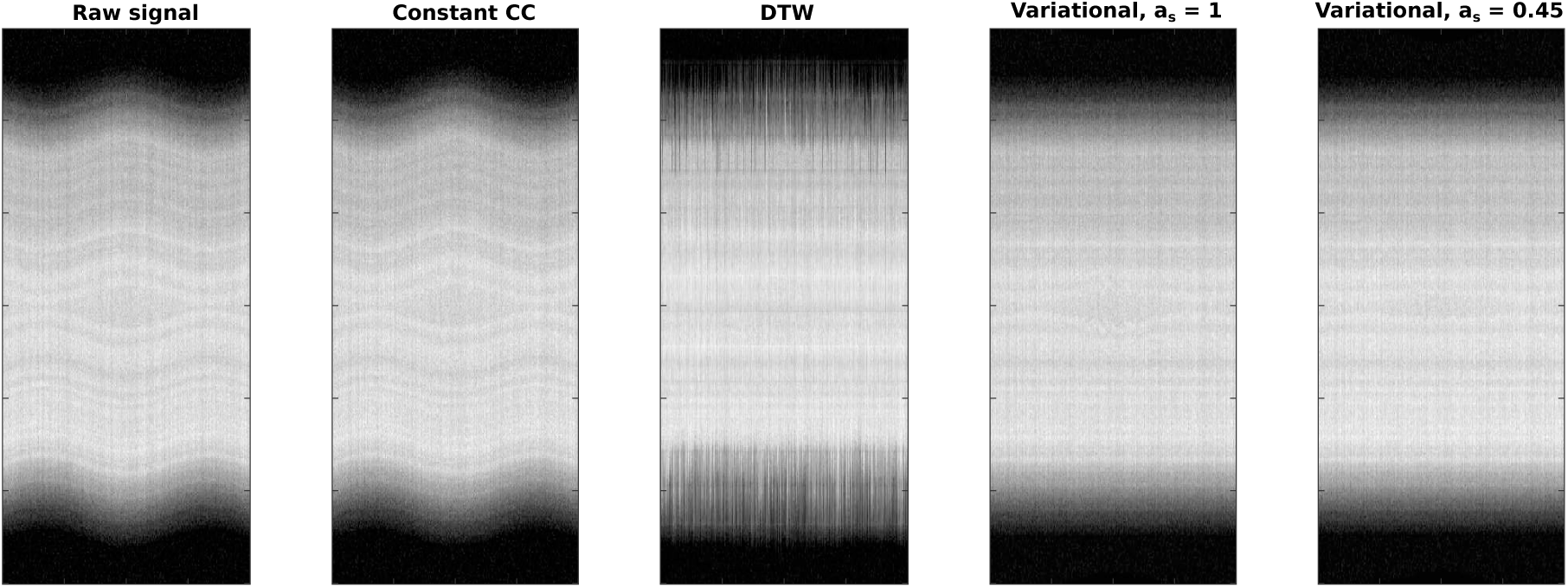
Qualitative registration performance on the first synthetic line scan recording. Left to right is the *raw signal*, removal of constant shifts with *cross correlation* (Constant CC), *correlation optimized warping* (DTW) and *variational alignment* with different parameter choices *a*_*s*_. Due to absence of constant shifts, constant CC cannot improve the input data. DTW contains many outliers and misalignments while variational alignment with *a*_*s*_ = 0.45 can almost perfectly align the sequence.

#### Alignment of Random Noise with structured Reference

We observe that our method tends to overfit noisy data if we do not apply pre-smoothing. This results in clustering of vertical traces (see figure 4 and 5), that is, if the SNR is low in a single line, features in the noise are aligned with respect to features in the reference. When we provide a structured pattern such as a sine as reference, overfitting can reproduce the pattern in the signal average after alignment (see figure 4). Regarding the average of an aligned matrix with the average profile as reference, overfitting seems to accentuate features in the average signal after alignment.

**Figure 4:**
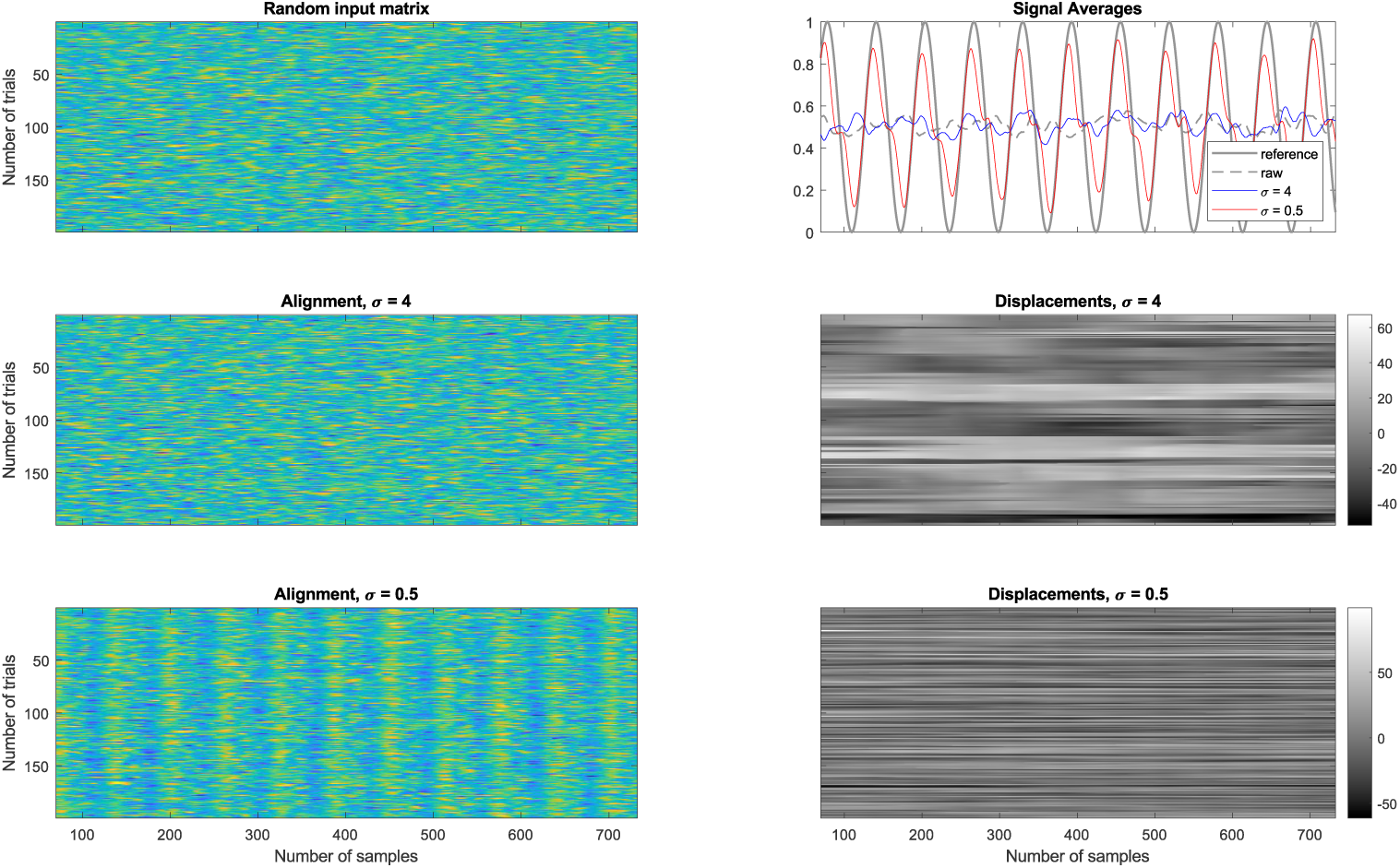
Alignment of random signals: We have aligned a matrix containing random, temporally lowpass filtered values with a sine wave as reference. It demonstrates the importance of integrating temporal coherence. When pre-filtering with *σ* = 4, the displacement field is relatively smooth across trials. With *σ* = 0.5 our method overfits and can reconstruct a reference that is not present in the data.

**Figure 5:**
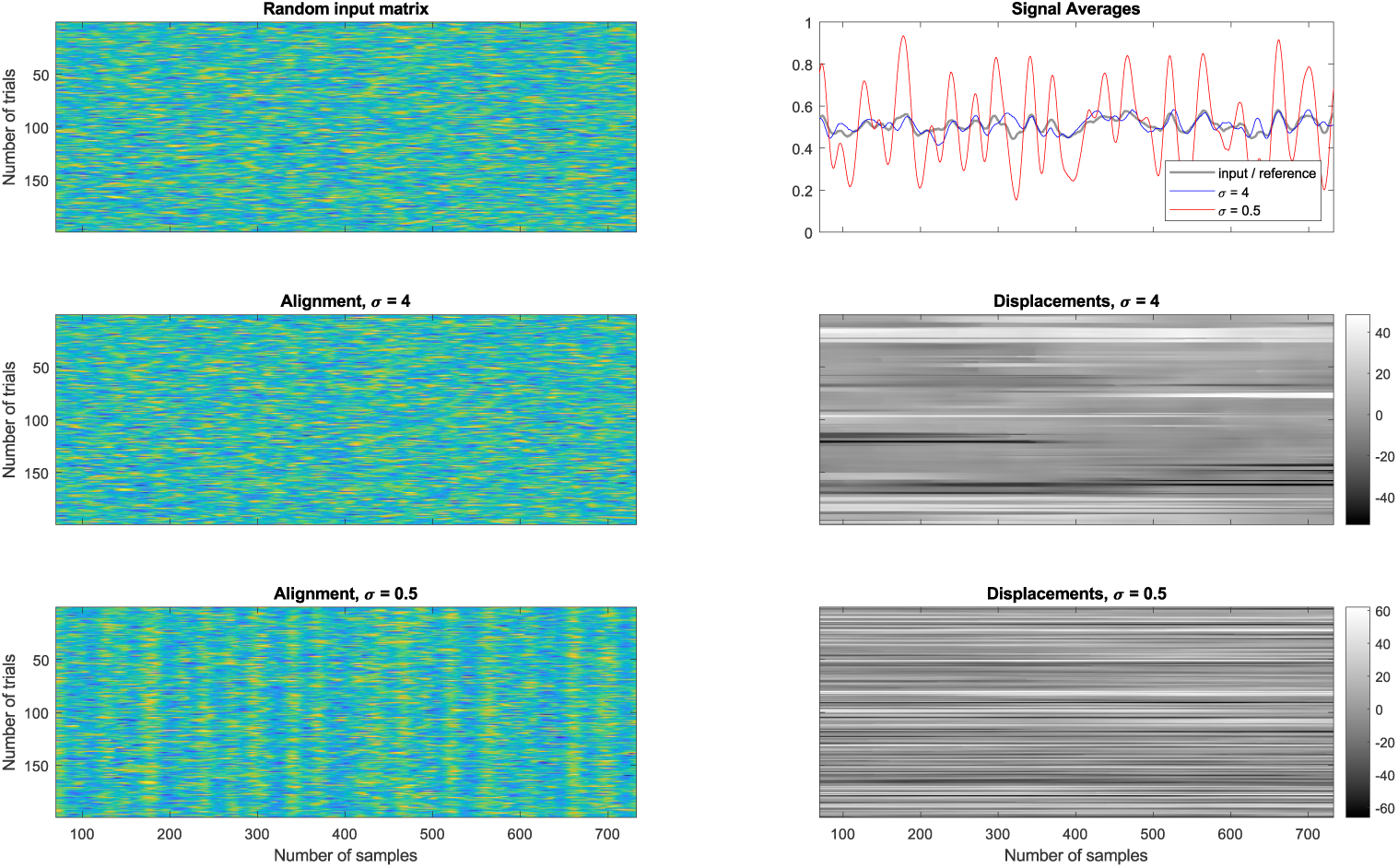
Alignment of random signals: When using the average line profile as reference, we observe a clustering effect when using a small filter size.

In our method, overfitting is regularized with the size of our filter from equation 4 across trials and the size of 1D filter along trials. Additionally, the subquadratic data term reduces the influence of outliers. The underlying assumption for the filtering across trials is a self-similarity in terms of a row coherence with respect to the provided permutation mapping *π*(*j*). Our anisotropic filter enhances alignment relevant regularities, while suppressing fluctuations that can be attributed to noise. Therefore, depending on the application, other denoising methods can be applied as well (e.g. compare [18], [19]). It is beyond the scope of this work to present a universal solution to avoid overfitting with this method. However, a general good practice is to adjust parameters, based on SNR. Problems where we generally deal with higher SNRs, or where temporal filtering can increase the SNR produce good results without much inter-trial filtering (see figure 8). Problems with inherently low signal to noise ratio are prone to overfitting and require pre-filtering, e.g., compare [18] and figure 10.

**Figure 6:**
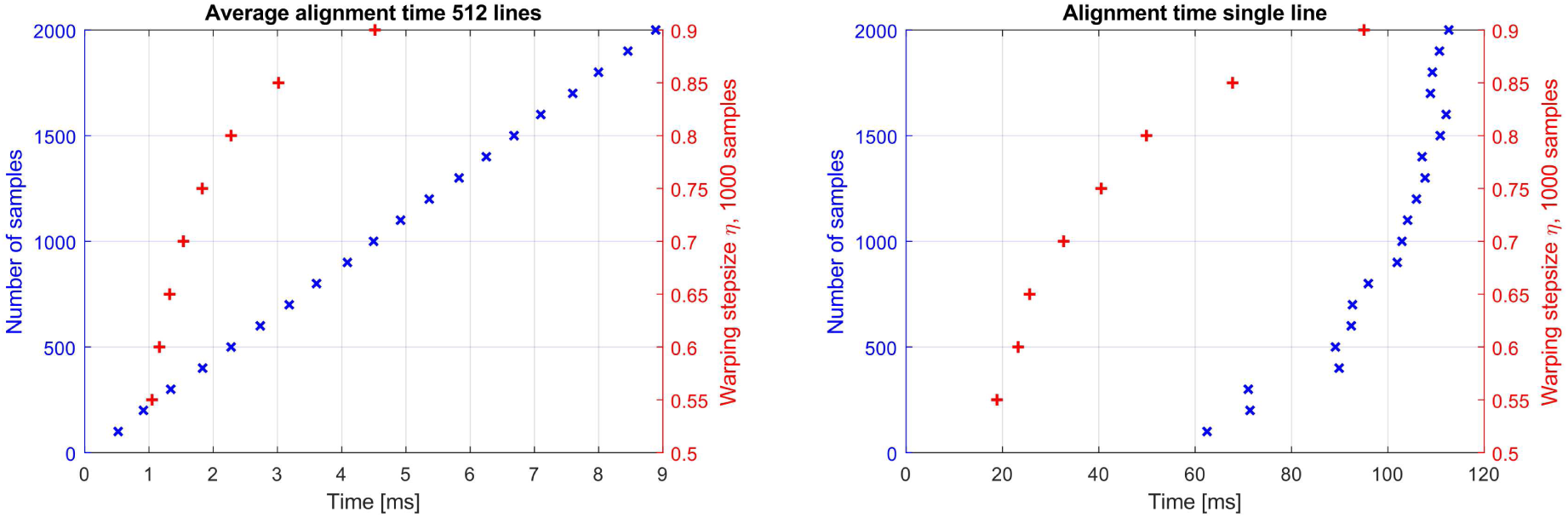
Alignment speed of our method per line with respect to varying number of samples and warping stepsize. We used 20 iterations and *η* = 0.9 for the varying sample size and 1000 samples for varying *η*. Left is the average time per line when using a multitrial matrix as input and right is the performance for the alignment of an isolated line.

**Figure 7:**
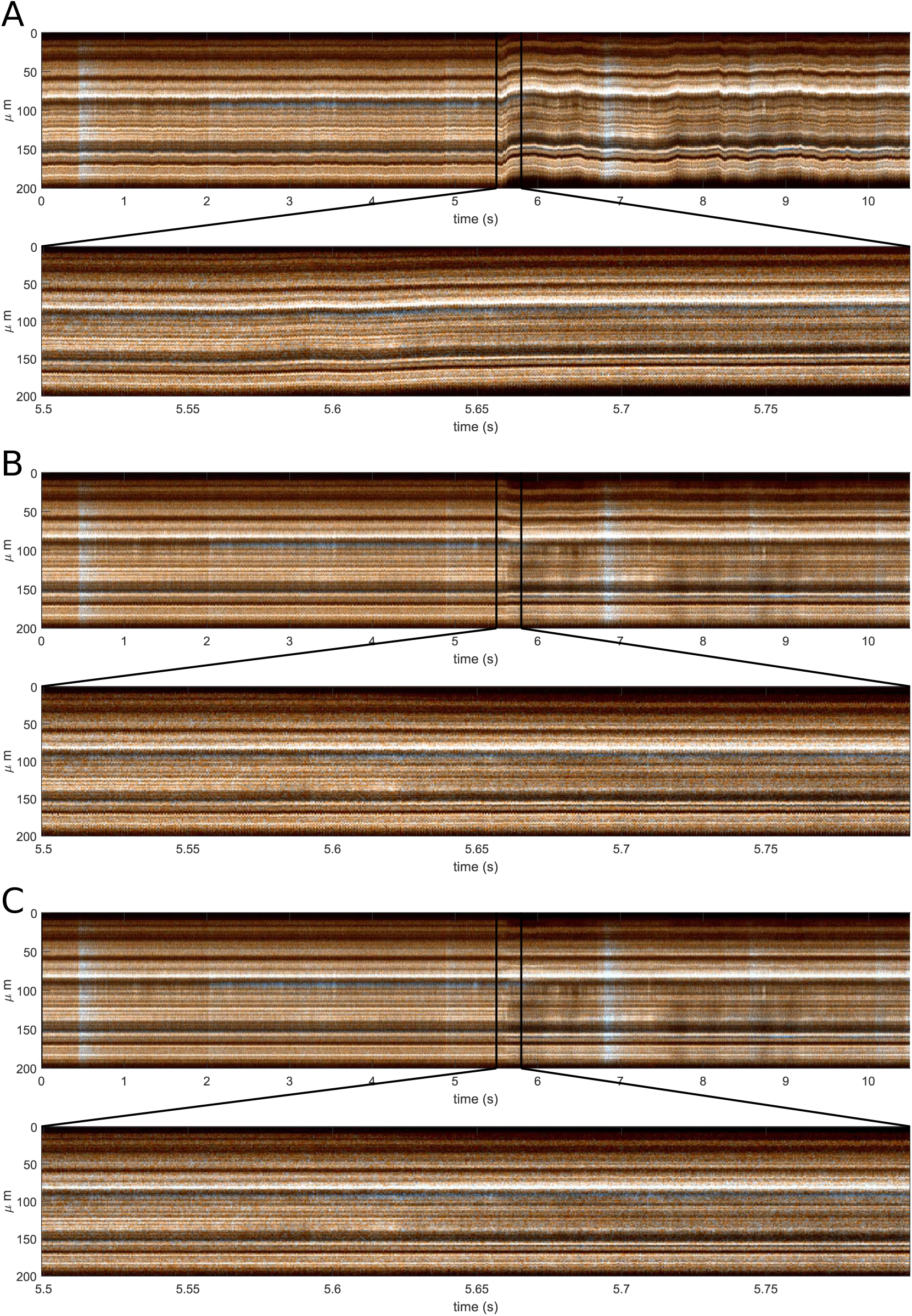
2-photon imaging line scan recording *without alignment* (A), with *cross correlation alignment* (B) and with *variational alignment* (C). The cross correlation approach fails to remove high frequency motion. During the strong motion artifact (5.5 s), removal of constant displacements cannot account for the converging structures. This results in an average temporal standard deviation of 241.2 for (A), 190.6 for (B) and 179.7 for (C). The recording is taken from Roome et al. [9] and used with the authors permission.

**Figure 8:**
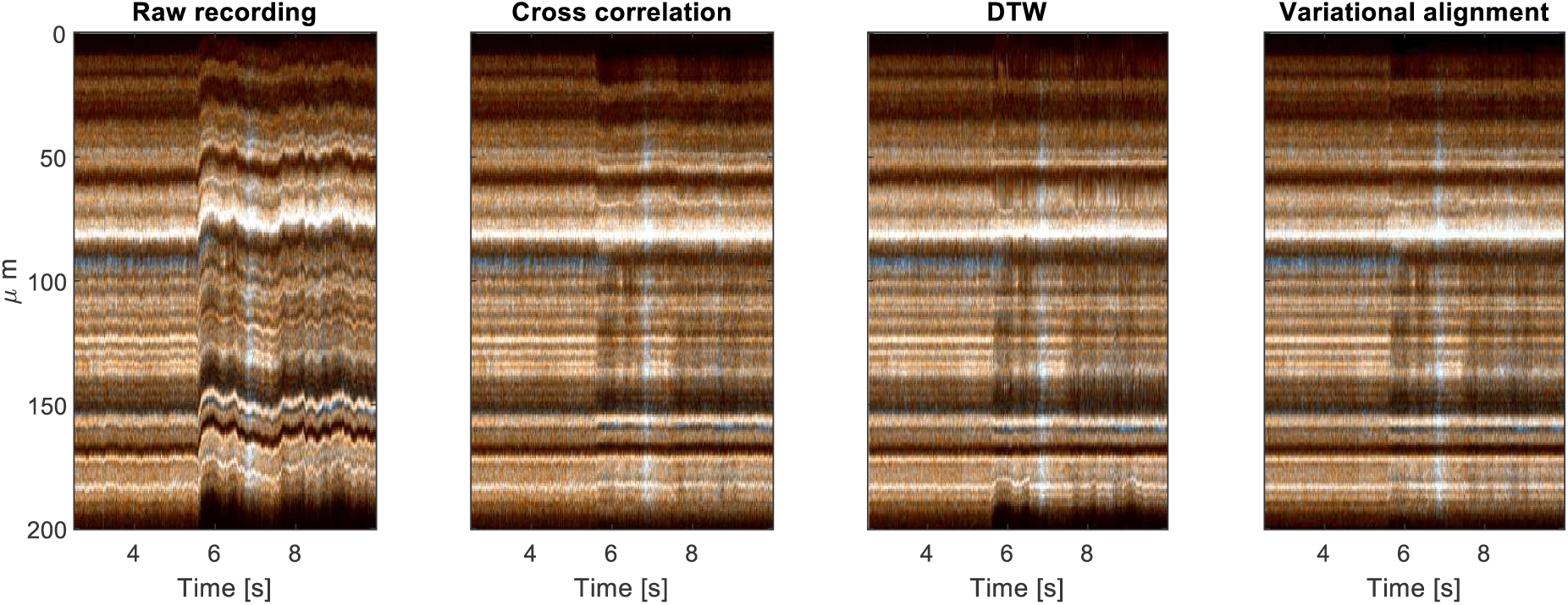
Section from the 2-photon imaging line scan recording from left to right *without alignment*, with *cross correlation alignment*, with *correlation optimized warping alignment* and with *variational alignment*.

**Figure 9:**
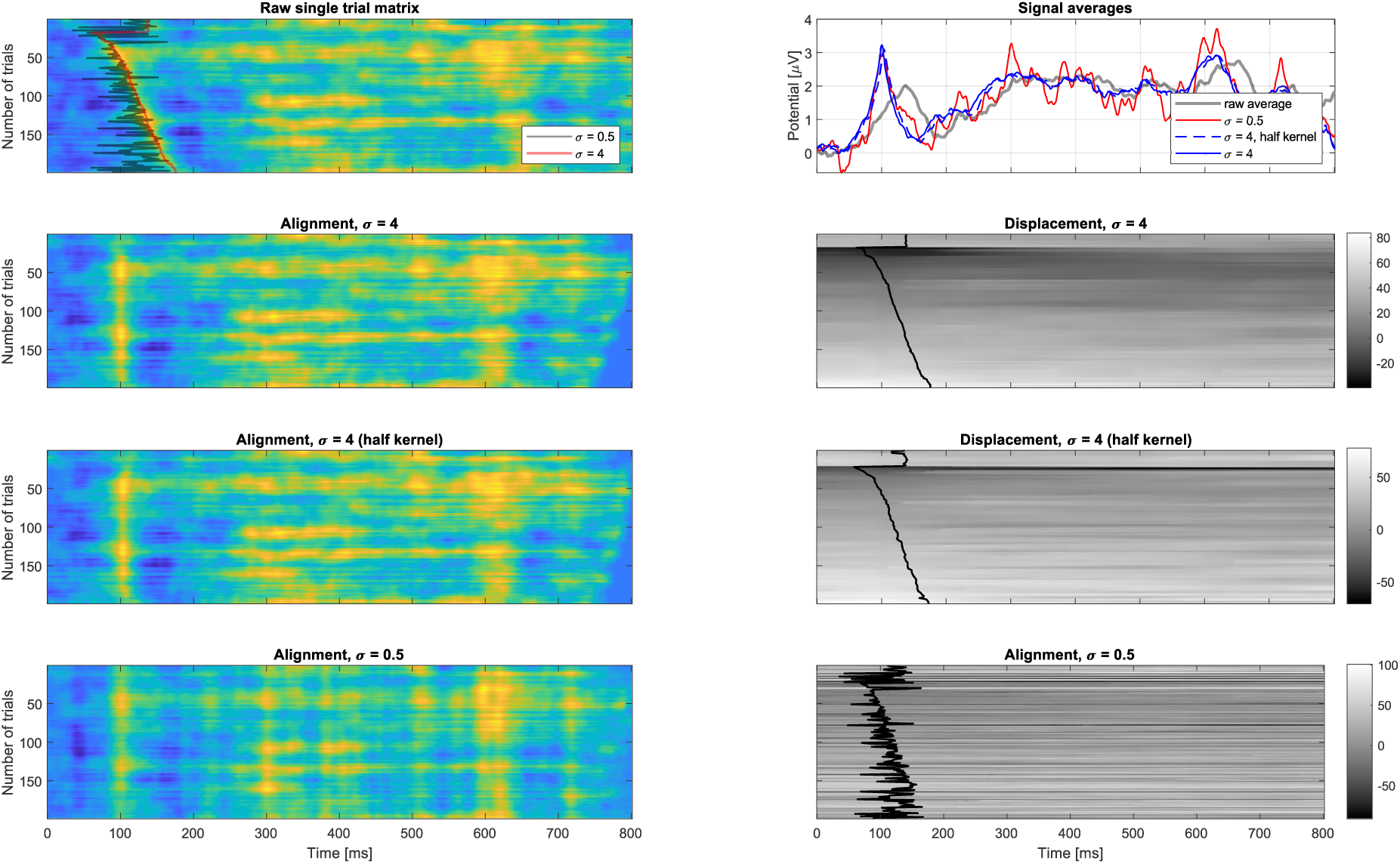
Alignment of an ERP recording sorted by subject’s response time, see [11]. Using only past trials results in a similar performance as the full kernel. However, a low level of temporal integration results in strong clustering of the signal. All ERP maps where filtered with *σ* = 4 for better visualization. The estimated trace is marked in black in the displacement maps on the right.

**Figure 10:**
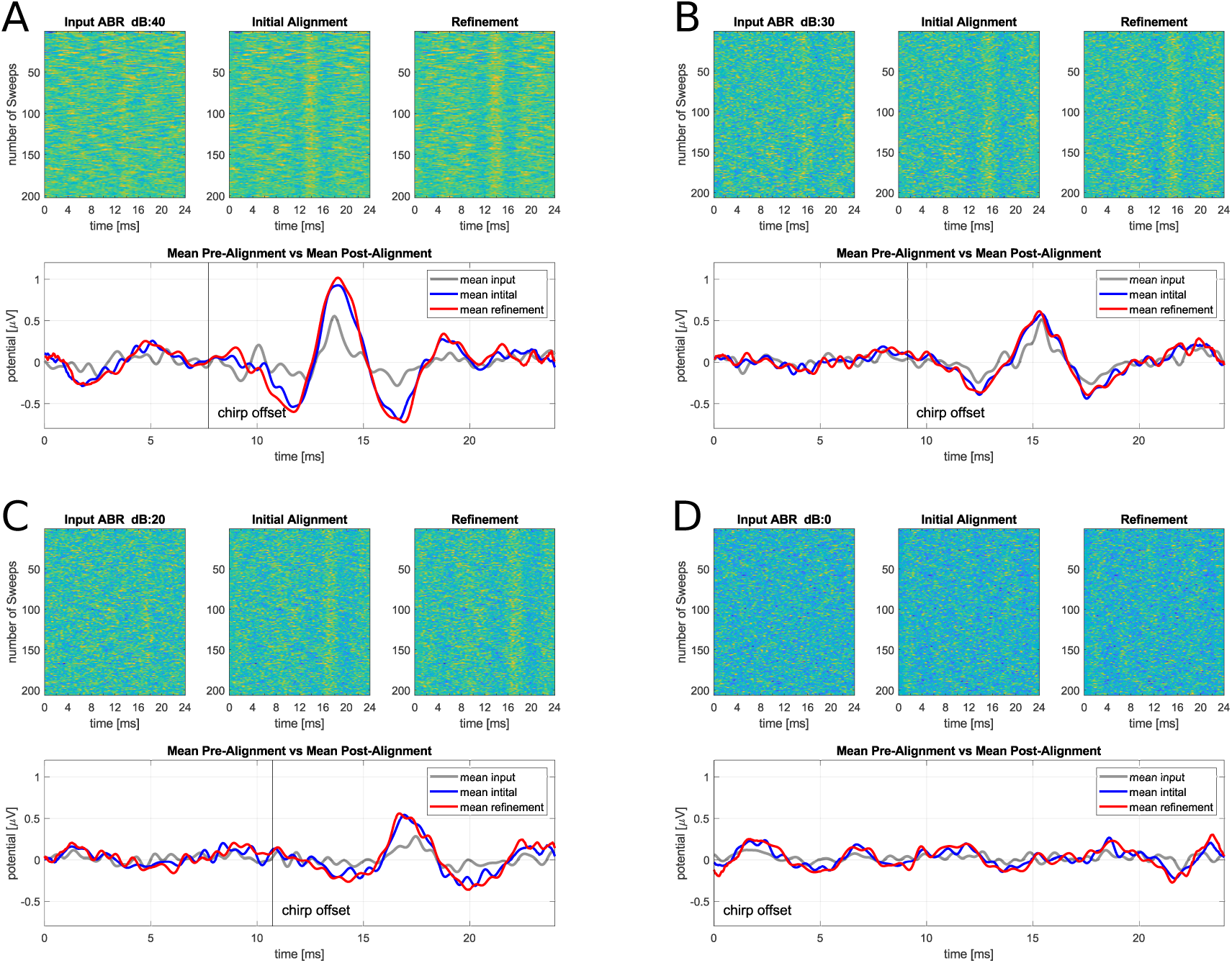
Application of our method to *auditory brainstem responses* (ABRs) data. In the average response, the diagnostically most relevant wave V is easier to recognize and its amplitude is enhanced after alignment as compared to classical averaging without alignment. However, we can also observe the clustering effects described in section 3.1 for lower stimulation amplitudes. Stimulation is done with 40 dB dB (A), 30 dB (B), 20 dB (C) and (D) is spontaneous activity. The refinement step is registration with the average of the matrix after initial alignment as reference.

We recommend to filter spatio-temporally such that the average SNR of signal and reference / average is positive. This way, the alignment only compensates visible displacements in the data. Also traces that appear after alignment without any visible correlates in the input data should be interpreted with caution and according to the respective application.

#### Real-Time Applications

Our method is fast enough to run in real-time on consumer grade hardware for different use cases. Given a varying sample size *n*, we benchmark the processing time of our MATLAB implementation on 512 × *n* matrices as well as on single lines with *n* samples and with a fixed number of 1000 samples for varying warping stepsizes *η* with 20 iterations (see figure 6). The alignment of the matrices took less than a few seconds with the average time per line in the order of milliseconds and no more than 120 ms for an isolated registration of a single line. The machine used for the estimation was a desktop computer with an AMD Ryzen 3900X processor (3.8 GHz, 12 cores, 24 logical processors) and 64 GB of memory. The alignment of an isolated line can be applied in the context of EPs and ERPs (see section 3.3), where we experiment with inherent delays of more than 100 ms in-between events. The registration of a full matrix might be applied to the online alignment of line scan data with block-wise acquisition and processing / display of line scan data.

For the alignment of short segments with *η* = 0.9 or for 1000 samples with small warping stepsize, our method performs with up to 1000 Hz. When aligning ERPs (see section 3.3), *η* = 0.9 with 1000 samples allows the compensation of a 2 s segment in 4.5 ms given enough room for other processing steps. In their preliminary paper, Deriso et al. [16] report an average speed of 0.25 s for a 1000 sample segment. Even though we cannot directly compare the results due to different hardware architectures, the sequential version of our method can perform around two magnitudes faster on average for multiple lines and depending on the choice of *η* can be one magnitude faster for the alignment of an isolated line.

When comparing our method with popular DTW approaches, on average our system needs 4.6 ms for 1000 samples and default parameters in MATLAB. On our system, the python package `fastdtw` takes around 229.6 ms with default parameters and 6 s with radius of 50. We constitute the slower alignment of isolated lines mainly to the characteristics of our MATLAB implementation: On the one hand, for multitrial input, filtering and resizing methods benefit from parallelized implementations in MATLAB. On the other hand, MATLAB overhead for calling external functions amortizes for larger matrices and is much more dominant for the alignment of individual lines. Also given the fact that the type of solver we use can be implemented on graphics cards and can easily be parallelized [7], we expect the performance of an optimized implementation of our method for alignment of a single line to be close to the reported average performance.

### 3.2 Line Scan Data

We apply our method to a 2-photon line scan recording from Roome et al. [9] of Purkinje neuron dendrites in the cerebellum of awake mice.

The recording contains two simultaneously recorded line scan channels, the first reporting calcium changes, the second voltage changes, and was recorded at 2 kHz. Imaging artifacts occurred because of animal movement and are composed of different components, including a slight convergence of the structures after movement onset, see figure 8 and 7 around 5.5 s. In the same way as for section 3.1, the performance of our method is compared with cross correlation based alignment of constant displacements and correlation optimized warping.

Optical imaging recordings can contain multiple channels [9] and sometimes contain a structural as well as a functional channel. We register on both channels simultaneously and weigh each channel by structure content.

For the alignment of the data in figure 7 and 8 our method took on average 2.2 ms per line with 100 iterations and with 20 iterations only 0.6 ms. From table 2 we see that there is not a large increase in performance for 100 compared to 20 iterations. Thus, state of the art results for a line scan of width 270 can be achieved at a little more than 1.6 kHz, which conforms with our results from the previous section.

**Table 2:**
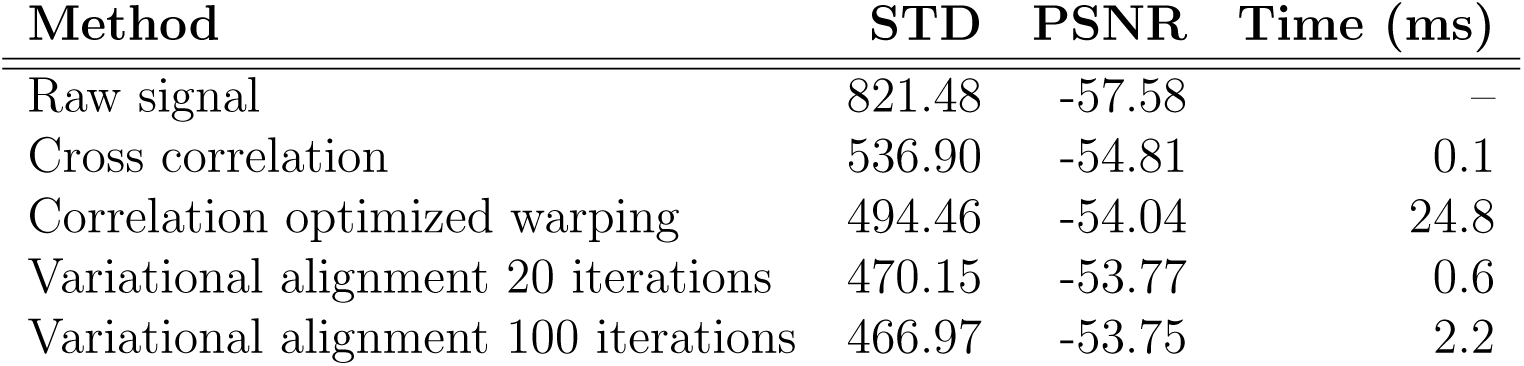
Performance of our method on the 2-photon imaging line scan data (unnormalized) in comparison with cross correlation based constant alignment (no subsample refinement) and correlation optimized warping. Lower values for the average *standard deviation* (STD) and higher values for the average *peak signal to noise ratio* (PSNR) are better.

However, difficult motion artifacts in two-photon imaging data as seen from 5.5 s onwards can coincide with shifts orthogonally to the imaging plane. This way, structures can completely disappear and concentration gradients of dyes can be misinterpreted as functional activity. Therefore, such points in time are usually detected and discarded systematically in evaluations. With an improvement in registration accuracy, it might be possible to use the previously discarded data for analysis. Importantly, the improved registration accuracy does not come at the cost of much higher computation time. Therefore, we see our method as a superior alternative to alignment approaches that estimate constant displacements.

### 3.3 ERP Single Trial Analysis

Analysis of 2D matrix representations of ERP in EEG recordings has been suggested for the analysis of trial to trial variances in critical wave morphologies, (e.g. see [11]). Inter-trial and inter-subject variances not only distort signal and grand averages, but also cannot be analyzed after averaging – even though they can contain different cues on dynamics during experiments, in particular, when analyzing endogenous processes and exogenous parameter variations. Because of the low SNR in individual trials, single trial analysis requires denoising strategies, which, due to the possibility of 2D representation of the data, are often motivated by image denoising techniques [11], [18].

Knowledge of the displacement or latency jitter in ERP matrices on a per trial basis has the potential to greatly improve ERP single trial analysis: In the context of denoising, aligning signals with respect to their morphology allows smoothing vertically to the time dimension with linear filters. Compared to anisotropic filtering approaches, the direction of inter-trial changes does not need to be estimated. Filtering methods based on non-local means estimation such as [11] produce state of the art results for two dimensional ERP denoising. However, they do not aim to recover the implicitly estimated spatial relation of the self-similarity. Also, the displacement maps in our method allow the analysis of the estimated traces (see section 3.1 and figure 9), while boosting important features in the signal average after alignment^1^. The same holds true for EP alignment: We apply our method to auditory brainstem responses (ABR) collected in Corona-Strauss et al. [20]. We observe, that the diagnostically most relevant wave V [21] is easier to recognize and its amplitude is enhanced after alignment as compared to classical averaging without alignment, see figure 10. This has the potential to reduce the required number of trials for analysis.

## 4 Discussion

In this work, we have presented a novel variational method for the alignment and estimation of non-flat and non-parametric displacements of 1D data with applications in neuroimaging. Our method produces state of the art results for the alignment of 1D signals and is fast enough to enable real-time applications on consumer grade hardware. The method is implemented as an accessible MATLAB toolbox and publicly available on GitHub^2^. Our continuous formulation is motivated by 2D OF estimation and thereby builds on a rich framework of different regularizers and constancy assumptions as well as efficient optimization strategies. Our model uses a total variation data term and quadratic or linear smoothness term. It natively supports weighted multichannel data. We have demonstrated the performance of our approach on 2-photon line scan recordings as well as 2D single trial matrix representations of ERPs and EPs. The alignment of ERP matrices opens the way for improved analysis with reduced number of trials. Due to its high accuracy and fast implementation forms an alternative to established methods such as DTW and constant template matching. In future work it can to be additionally evaluated against methods currently under development such as Deriso et al. [16] (unpublished work).

For line scan recordings, our method is an alternative to constant displacement estimation with higher accuracy, only slightly less computation time and the ability to compensate non-flat displacements. In addition, we believe that this method could find many further applications for the alignment of line scan data beyond the field of neuroimaging.

## 5 Acknowledgements

This work was partially funded by the Japan Society for the Promotion of Science (JSPS) with the Summer Program 2019 and partially conducted at the *Optical Neuroimaging Unit* under Prof. Bernd Kuhn at the Okinawa Institute of Science and Technology Graduate University.

## Footnotes

1 For an in depth analysis of the application of our method to ERP data, we refer the reader to Thinnes et al. [18].

2 https://github.com/phflot/variational_aligner

